# Characterization of a Novel *FKS1* Mutation in *Candida lusitaniae* Shows a Potential Critical Role for *MKC1* in Echinocandin Resistance

**DOI:** 10.1101/2025.04.08.647726

**Authors:** Isabelle Accoceberry, Maxime Lefranc, Camille Meunier, Fabienne Quilès, Milena Kordalewska, Nicolas Biteau, David S Perlin, Sofiane El-Kirat-Chatel, Thierry Noël

## Abstract

Caspofungin is an echinocandin antifungal that inhibits glucan synthesis in the fungal cell wall. A *Candida parapsil*osis bloodstream isolate resistant to echinocandins was recovered from a patient who had undergone allogeneic hematopoietic stem cell transplantation. The *FKS1* gene, encoding the target glucan synthase, contained a heterozygous mutation resulting in an I1380T amino acid change, in addition to the naturally occurring P660A polymorphism. When expressed at the equivalent position in the Fks1p protein of *C. lusitaniae*, P642A and I1359T, alone and in combination, led to 6-, 12-, and ≥256-fold increases in the minimal inhibitory concentration (MIC) of caspofungin, respectively. The caspofungin concentration needed to inhibit 50% of glucan synthase activity was increased 3-, 37-, and 270-fold, respectively. At high drug concentrations, and also in drug-free medium, infrared spectroscopy revealed a decrease in β-glucan content and an increase in chitin in the cell wall of the I1359T Fks1p mutants. Atomic force microscopy showed cell wall damage and cell swelling in both susceptible and resistant strains under caspofungin exposure. Analysis of susceptibility to cell-wall stressors and key factors in cell wall integrity (CWI) and high-osmolarity glycerol (HOG) pathways showed that all strains activated these pathways under caspofungin stress. In the I1359T Fks1p mutants, Mkc1p was constitutively activated even without caspofungin. Deletion of *MKC1* restored caspofungin susceptibility, indicating that activation of the CWI pathway is a key molecular determinant of resistance *in vitro* to caspofungin in these mutants.

## INTRODUCTION

Invasive candidiasis is a common hospital-acquired infection with a high crude mortality rate. Although *Candida albicans* is the predominant pathogenic yeast species, non*-albicans Candida* species, some of which have reduced susceptibility to antifungals, account for more than half of all cases in some geographical areas (1, 2). *Candida parapsilosis* generally ranks second or third among the species most often isolated from invasive infectious episodes, especially in neonates and critically ill patients in intensive care units (3). This species is diploid and belongs to the CTG clade of yeasts, which decode the CUG codon as serine instead of leucine (4). Numerous genetic traits contribute to *C. parapsilosis* pathogenesis, such as adherence proteins, secreted enzymes, and the formation of biofilm on human skin and implanted medical devices, which may serve as a starting point for invasive candidiasis (5, 6).

*C. parapsilosis* has reduced *in vitro* susceptibility to echinocandins, which are the first-line treatment for patients with invasive candidiasis (7). Echinocandins inhibit the 1,3-β-glucan synthase responsible for the biosynthesis of the major glucan component of the yeast cell wall (8). The reduced susceptibility of *C. parapsilosis* stems from the species-specific amino acid polymorphism P660A in the glucan synthase (9). We have reported that expression of a *FKS1* allele bearing the P660A polymorphism at the equivalent position in *Candida lusitaniae* conferred a six-fold increase in the caspofungin MIC (10). In *C. albicans*, the main mechanism of echinocandin resistance involves non-synonymous mutations in two hotspot regions, HS1 and HS2, of the *FKS1* gene (11), altering drug-target interactions (12). In *C. parapsilosis*, with the exceptions of two recently reported HS1 mutations (13, 14), the *FKS* polymorphisms described to date were located outside the HS1 and HS2 regions (15), and their role in resistance was not functionally demonstrated *in vitro*. Additionally, cellular stress response pathways involving calcineurin, high-osmolarity glycerol (HOG), and protein kinase C (PKC), can contribute to increased chitin synthesis (16), cell wall remodeling (17), and tolerance to echinocandins (18).

In this work, we identified a new mutation in a *FKS1* allele of an echinocandin-resistant *C. parapsilosis* clinical isolate responsible for invasive candidiasis. In order to study the real impact of this mutation on echinocandin resistance and its biological effects, we chose to express it in a genetic context different from the one in which it was selected in the clinic. To do this, we used a strain of the haploid species *C. lusitaniae*, genetically modified for the expression of full-length *FKS1* alleles in an isogenic context (10). We then determined the caspofungin kinetic inhibition parameters of the variants of 1,3-β-glucan synthase thus obtained. Morphological changes induced by caspofungin were analyzed using atomic force microscopy (AFM), which enables nanometer-level imaging of the properties of yeast-cell surfaces exposed to external stresses (19–22). Caspofungin-induced morphological changes were correlated with variations in the cell wall biochemical composition using infrared spectroscopy in attenuated total reflection mode (IR-ATR), a non-invasive technique enabling qualitative and quantitative analysis of yeast cell wall composition (23–25). Although the glucan synthase of *C. lusitaniae* expressing this new mutation along with the genetic polymorphism specific to *C. parapsilosis* had decreased binding capacity for caspofungin, glucan synthesis seemed to be altered even in the absence of drug. This resulted in a reduced 1,3-β-glucan content in the cell wall, increased chitin synthesis, constitutive activation of the Cell Wall Integrity (CWI) rescue pathway, and global loss of fitness during interactions with murine macrophages *in vitro*.

## RESULTS

### Isolation of a *Candida parapsilosis* strain resistant to echinocandins

This clinical case involved a 35-year-old woman who underwent allogeneic hematopoietic stem-cell transplantation (HSCT) for an idiopathic medullary aplasia with paroxysmal nocturnal hemoglobinuria positivity. The patient remained neutropenic over the subsequent four months and was empirically treated with several courses of broad-spectrum antibiotics and caspofungin for septic episodes with a high level of C-reactive protein (230 mg/L). On Day 90 of HSCT, while the patient was on caspofungin (120 mg/daily), yeast was isolated from a blood sample and identified by MALDI-TOF mass spectrometry as *C. parapsilosis* (CPAR). The MICs for antifungal agents measured by Etest at 48 h were caspofungin, 32 μg/mL; micafungin, 32 μg/mL; anidulafungin, 32μg/mL (Supplemental Data S1); fluconazole, 0.094 μg/mL; voriconazole, 0.012 μg/mL; 5-fluorocytosine, 0.012 μg/mL; and amphotericin B, 0.19 μg/mL. Because the CPAR isolate was resistant to echinocandins, nucleotide sequencing of the entire *FKS1* gene was performed. Comparison with the sequence of the *FKS1* gene of *C. albicans* SC5314 confirmed the presence of the naturally occurring homozygous polymorphism C1978G leading to the P660A (proline replaced by alanine) amino acid substitution in the HS1 region (amino acids 652 to 660), which has been described in *C. parapsilosis* (9). Comparison with the sequence of the *FKS1* gene of the caspofungin-susceptible *C. parapsilosis* CBS 604 revealed the T4139C heterozygous mutation, leading to the substitution I1380T (isoleucine replaced by threonine) four amino acids downstream of the HS2 region (amino acids 1369 to 1376) of glucan synthase (Supplemental Data S2).

### Fks1p P660A and I1380T substitutions are necessary for high-level caspofungin resistance *in vitro*

To evaluate the contribution of each mutation of the CPAR *FKS1* gene to echinocandin resistance *in vitro*, the nucleotide changes responsible for the P660A and I1380T substitutions were introduced separately and in combination at their equivalent positions in the *FKS1* gene of *C. lusitaniae*, yielding Fks1p bearing the P642A and 1359T substitutions (Fig. 1A). The newly generated *FKS1* alleles were transformed and expressed in *C. lusitaniae* F1 *trp1*Δ3’, *ura3*Δ5’, a caspofungin-susceptible strain (MIC 0.125 µg/mL) that was engineered from the wild-type (WT) strain CBS 6936 for the chromosomal allelic replacement of the essential *FKS1* gene (10). The resulting transformants were tested for caspofungin susceptibility. Compared to the WT, introduction of P642A in Fks1p resulted in a six-fold increase in caspofungin MIC (0.75 μg/mL), as reported previously (10). The I1359T replacement increased the caspofungin MIC 12-fold (1.5 μg/mL), and the combination of P642A and I1359T led to a caspofungin MIC ≥ 32 μg/mL, at least 256-fold higher than the WT and comparable to the *C. parapsilosis* CPAR clinical isolate (Fig. 1B). The growth curves obtained in liquid RPMI confirmed the phenotype of resistance to caspofungin in the mutant strains and showed that expression of mutated Fks1p did not alter the growth capacity of the mutant strains in drug-free RPMI (Supplemental Data S3).

**Figure 1.**
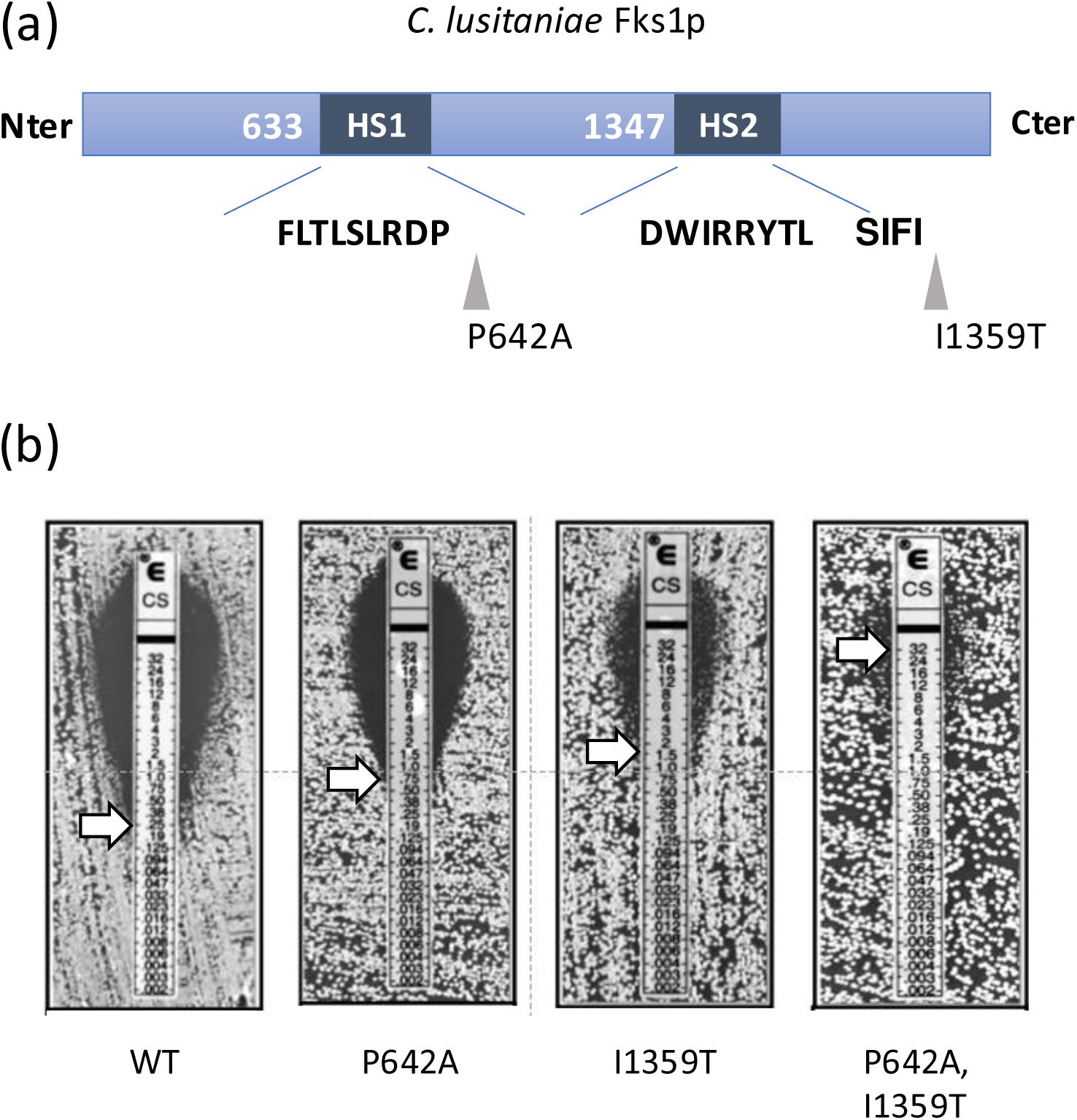
Caspofungin susceptibility of *C. lusitaniae* expressing different *FKSl* alleles. (a) Schematic position of amino acid substitutions introduced in *C. lusitaniae* Fks1 proteins equivalent to those identified in *C. parapsilosis* CPAR. (b) Etest determination of caspofungin susceptibility of the *C. lusitaniae* CBS 6936 strain expressing a wild type (WT) Fks1 protein and of the *C. lusitaniae* strains expressing Fks1 proteins containing P642A, 11359T, and both P642A, 11359T amino acid substitutions. For each *FKSl* allele expressed, caspofungin M1C was determined for six independent clones after having controlled at the molecular level that they had correctly integrated the SNP of interest. The M1C is indicated by a white arrow.

### High caspofungin concentrations are necessary to inhibit the activity of glucan synthase bearing the P642A and I1359T substitutions

To assess the direct inhibition of caspofungin on glucan synthase, the half-maximal inhibitory concentration (IC_50_) values were determined for 1,3-β-glucan synthases expressed by the *FKS1* WT and mutant strains of *C. lusitaniae* in polymerization assays using [^3^H]UDPG as the substrate (11). The inhibition curve for caspofungin against *C. lusitaniae* expressing WT enzyme displayed the typical pattern of 1,3-β-glucan synthase echinocandin susceptibility with a caspofungin IC_50_ of 13.36 ng/mL (Fig. 2). Decreased echinocandin susceptibility was observed with the *C*. *lusitaniae* expressing the P642A, I1359T, and P642A+I1359T mutant enzymes, with 3-, 37-, and 270-fold increases in echinocandin IC_50_ values, respectively, compared with the WT enzyme (Fig. 2).

**Figure 2.**
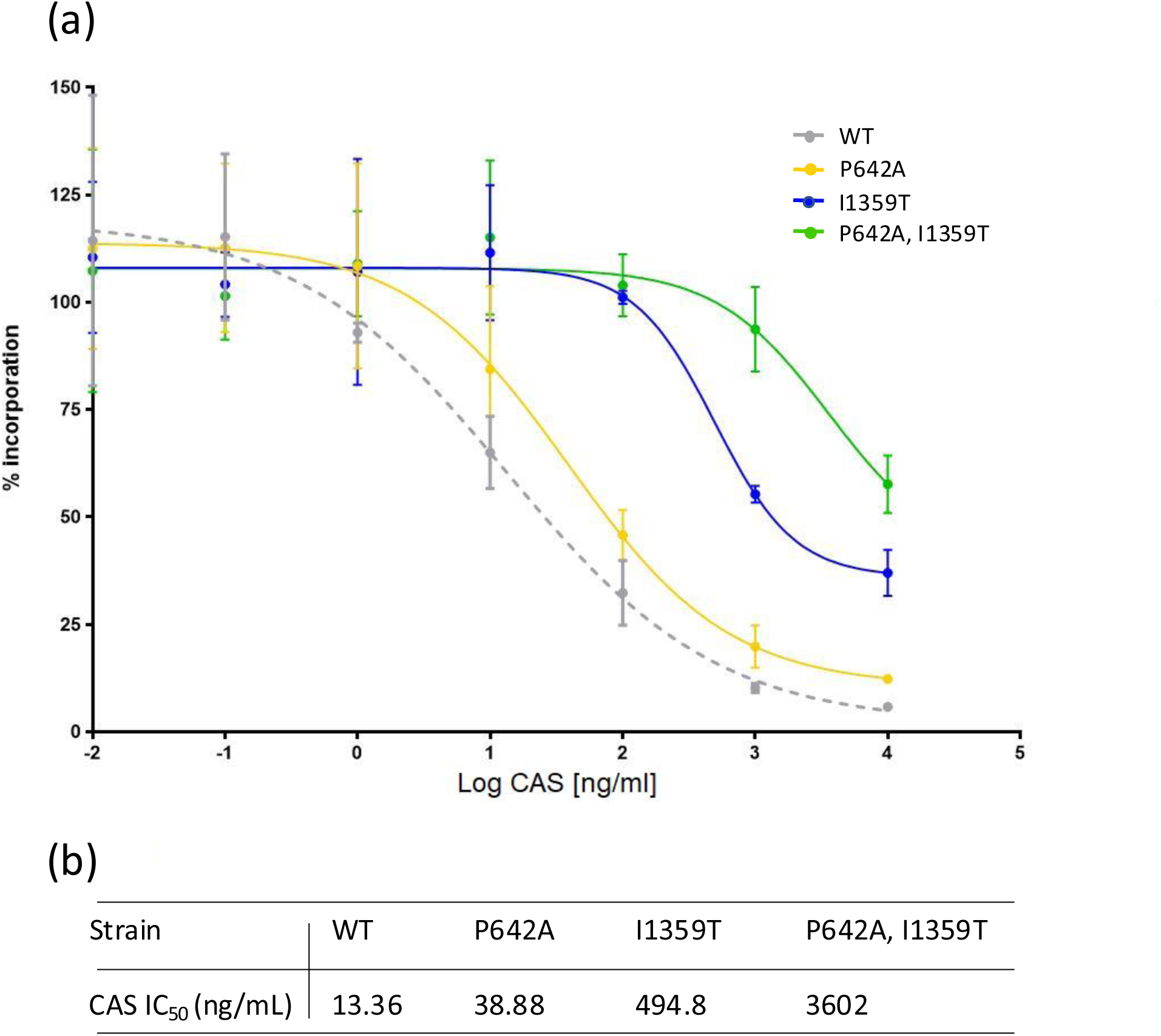
Caspofungin (CAS) inhibition profiles of enriched glucan synthase (GS) complexes from wild type and mutant *C. lusitaniae* strains. Relative GS activity to the wild type strain 6936 (WT) was assessed by the incorporation of [^3^H]glucose in glucan polymers. (a) Caspofungin titration curves for the *C. lusitaniae* WT strain, and strains expressing the Fks1 proteins with P642A, 11359T and P642A+11359T substitutions. (b) Caspofungin half maximal inhibitory concentration (1C_5O_) values for each *C. lusitaniae* strain.

### Introduction of I1359T in *C. lusitaniae* Fks1p leads to a reduced cell wall glucan content in cells exposed to high caspofungin concentration

We used the *FKS1* WT and mutant isogenic strains of *C. lusitaniae* to evaluate the effects of the *FKS1* variants on the composition of the cell wall by IR-ATR. IR-ATR spectra were recorded for yeast suspensions in PBS after 16 h of incubation in medium containing no drug and in medium containing a low (0.05 µg/mL; 0.4× WT MIC) or high (5 µg/mL; 40× WT MIC) caspofungin concentration (Fig. 3a). The integrated intensities of the bands are directly related to the concentrations of the biochemical compounds. In the region 1800 to 1200 cm^-1^, the spectra revealed mainly proteins (amides I and II at 1644 and 1546 cm^-1^, respectively) and phosphate compounds (1248 cm^-1^), which could correspond to nucleotide sugars, *i.e.*, the substrates of glucan and chitin synthases, and to phospholipids. Bands at 1377 and 1309 cm^-1^, assigned to chitin (26), were resolved upon treatment with caspofungin. In the region < 1200 cm^-1^, assigned mainly to C-O-C, C-OH, C-C and PO_2_ stretchings, the four *C. lusitaniae* strains exhibited main bands at 1150, 1081, 1064, 1028, and 1005 cm^-1^ (determined from the second derivative spectra, not shown). The bands at 1150, 1081, and 1005 cm^-1^ were assigned to 1,3-β-glucans (26) mixed with other glucans, possibly 1,4-β-glucans based on the band at 1064 cm^-1^ (27).

**Figure 3.**
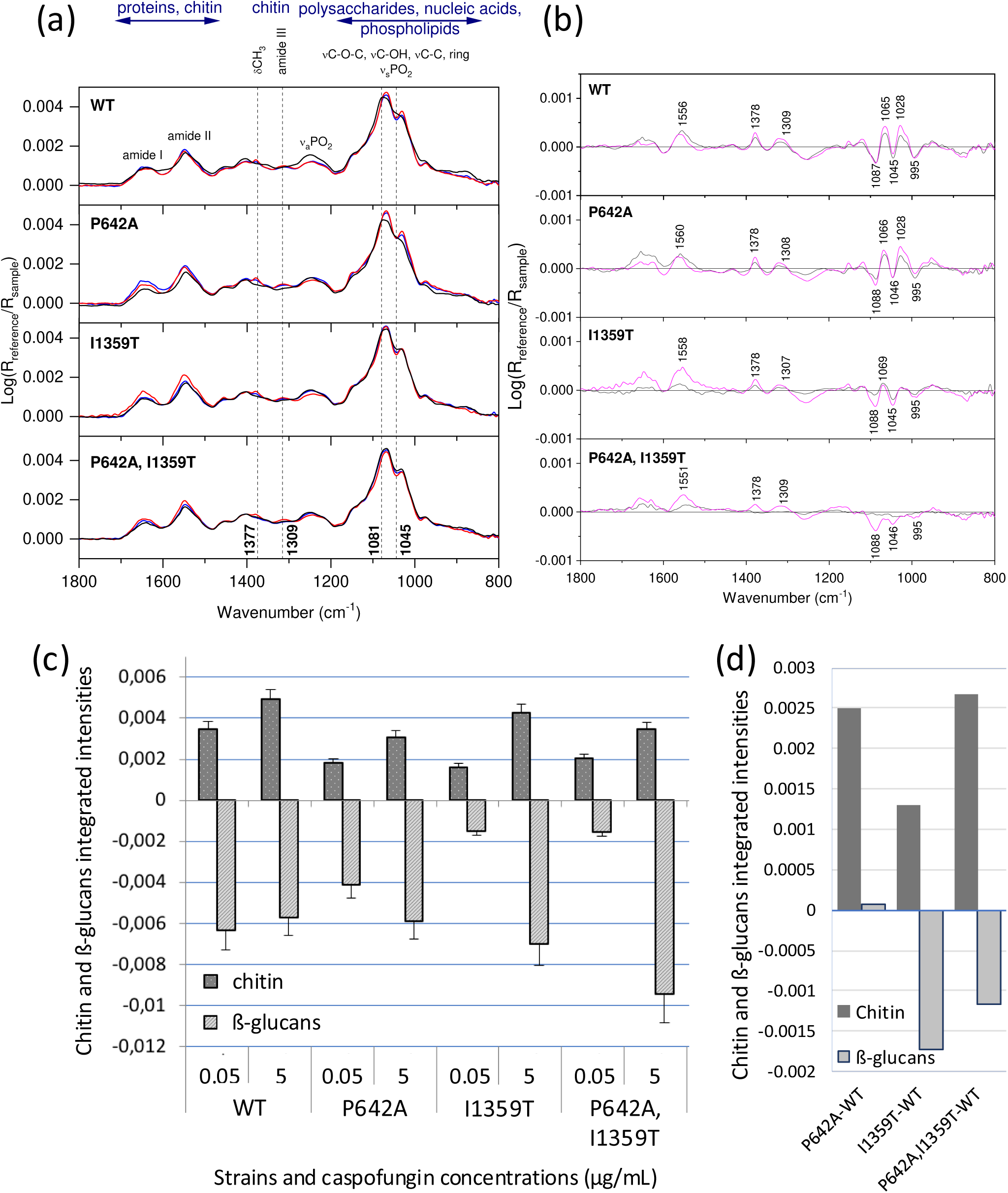
(a) Infrared spectra in ATR mode of yeasts in suspension in PBS, without treatment (black lines) and after treatment with caspofungin at 0.05 (blue lines) and 5 µg/mL (red lines) during 16 h. (b) Difference spectra between the spectra from untreated yeasts and those from yeasts treated with caspofungin at 0.05 (dark blue lines) and 5 µg/mL (pink lines). Key: ν, stretching; δ, bending; s, symmetric; a, antisymmetric. (c) Integrated intensities related to the variations in chitin and ß-glucans amounts in the cell wall of WT and *FKS1* mutant strains of *C. lusitaniae* after treatment with caspofungin. Integrated intensities were calculated from the difference spectra of Figure 3b as the values corresponding to the integrated intensities of CH2 bending bands from chitin, and C-C, C-O stretching band from ß-glucans (integrated regions: 1394-1361 and 1110-1077 cm-1, respectively). Y axis positive values: increase in chitin content in the cell wall relative to untreated cells. Y axis negative values: decrease in ß-glucan content in the cell wall relative to untreated cells. (d) Integrated intensities related to the variations in chitin and ß-glucans amounts in the cell wall of *FKS1* mutants grown in drug-free mediumafter subtracting the cell wall spectrumofWT grown in the same medium. Legends as in (c).

Figure 3b presents the spectra of yeast cells treated with caspofungin after subtracting the spectrum of cells grown in drug-free medium. The difference spectra of caspofungin-treated cells were characterized by positive bands assigned to chitin at 1556, 1378, and 1309 cm^-1^, and negative bands assigned to 1,3-β-glucans at 1087, 1045, and 995 cm^-1^ (26, 28). The increase in chitin synthesis and the decrease in 1,3-β-glucan content was caspofungin dose-dependent for all strains, except for ß-glucans in WT, and the degrees of increases and decreases differed according to the mutation in *FKS1* (Fig. 3c). At high drug concentration, the reduction in 1,3-β-glucan content was comparable in the WT and P642A, but the 1,3-β-glucan content decreased markedly in strains with the Fks1p I1359T mutation. Interestingly, the strain expressing Fks1p with both the P642A and I1359T substitutions had the greatest reduction in cell wall glucan content when exposed to high caspofungin concentration compared to untreated cells, suggesting marked alteration of glucan synthase activity.

This hypothesis appears to be corroborated by the comparison of the spectra of mutant cells grown in drug-free medium, after subtracting the spectrum of WT cells grown in the same medium (Fig 3d, derived from the spectra of Supplementary Data S4). The introduction of the I1359T mutation in Fks1p not only increased the chitin content in the cell wall, as with P642A, but also reduced the glucan content in the cell wall of both single and double mutant.

### Effects of the P642A and I1359T substitutions in Fks1p and of caspofungin treatment on cell topography

AFM was used to image WT and mutant *C. lusitaniae* cells in the absence of drug and after treatment with caspofungin at 0.05 and 5 µg/mL for 16 h. We first analyzed the effect of caspofungin on *C. lusitaniae* WT cells (Fig. 4a-c). Untreated WT cells had smooth and regular surfaces and some had elongated shapes typical of pseudohyphal growth (Fig. 4a). After caspofungin treatment, WT cells had rough surfaces with crests, hollows, and furrows (Fig. 4b-c). This change was accompanied by cell swelling and a wider cell size distribution (Fig. 5). Introduction of the P642A substitution into Fks1p resulted in yeast cells with a similar phenotype to WT cells, regardless of caspofungin treatment (Fig. 4d-f). Cells expressing P642A Fks1p are phenotypically as affected as WT cells by caspofungin, with a much more marked effect on cell size heterogeneity (Fig.5). Surprisingly, the I1359T substitution in Fks1p yielded some yeast cells with irregular rough surfaces even in the absence of drug (Fig. 4g), and the cells were larger and had a wider size distribution than WT and P642A cells (Fig. 5). After caspofungin treatment, I1359T mutant cells had a wrinkled surface (Fig. 4h-i). *C. lusitaniae* cells with the P642A and I1359T substitutions in Fks1p (Fig. 4j-l) exhibited morphological traits similar to the single I1359T mutant. These findings suggest that although both substitutions are required for full resistance to caspofungin *in vitro*, I1359T impacts Fks1p function and cell wall organization. This morphological analysis confirms that caspofungin treatment always affects cell surface topography regardless of its concentration or strain susceptibility, and reveals that the single amino acid substitution I1359T in Fks1p may be responsible for morphological changes even in cells not exposed to caspofungin stress.

**Figure 4.**
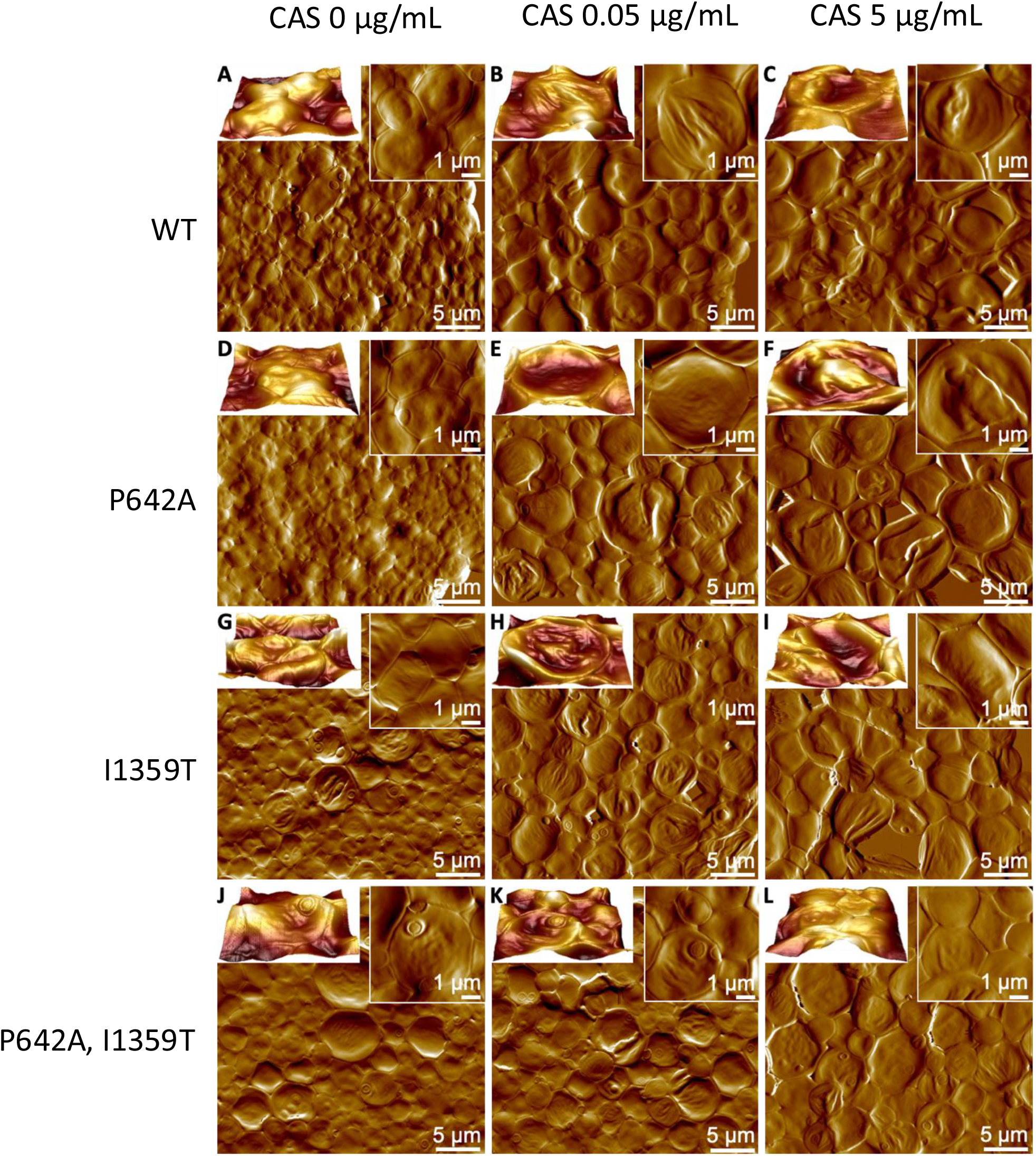
AFM images in air of *C. lusitaniae* WT, P642A, 11359T and P642A, 11359T cells without and with caspofungin treatment at 0.05 µg/mL and 5 µg/mL. Large scale deflection (error signal) images of WT (a, b, c), P642A (d, e, f), 11359T (g, h, i) and P642A, 11359T cells (j, k, l) without caspofungin treatment (a, d, g, j) and after treatment with 0.05 µg/mL of caspofungin (b, e, h, k) and 5 µg/mL of caspofungin (c, f, i, l). Top left insets are enlarged tridimensional views (merged height and deflection) showing the surface ultrastructure of individual cells in the corresponding condition. Top right insets are high resolution deflection images of cells in the corresponding condition.

**Figure 5.**
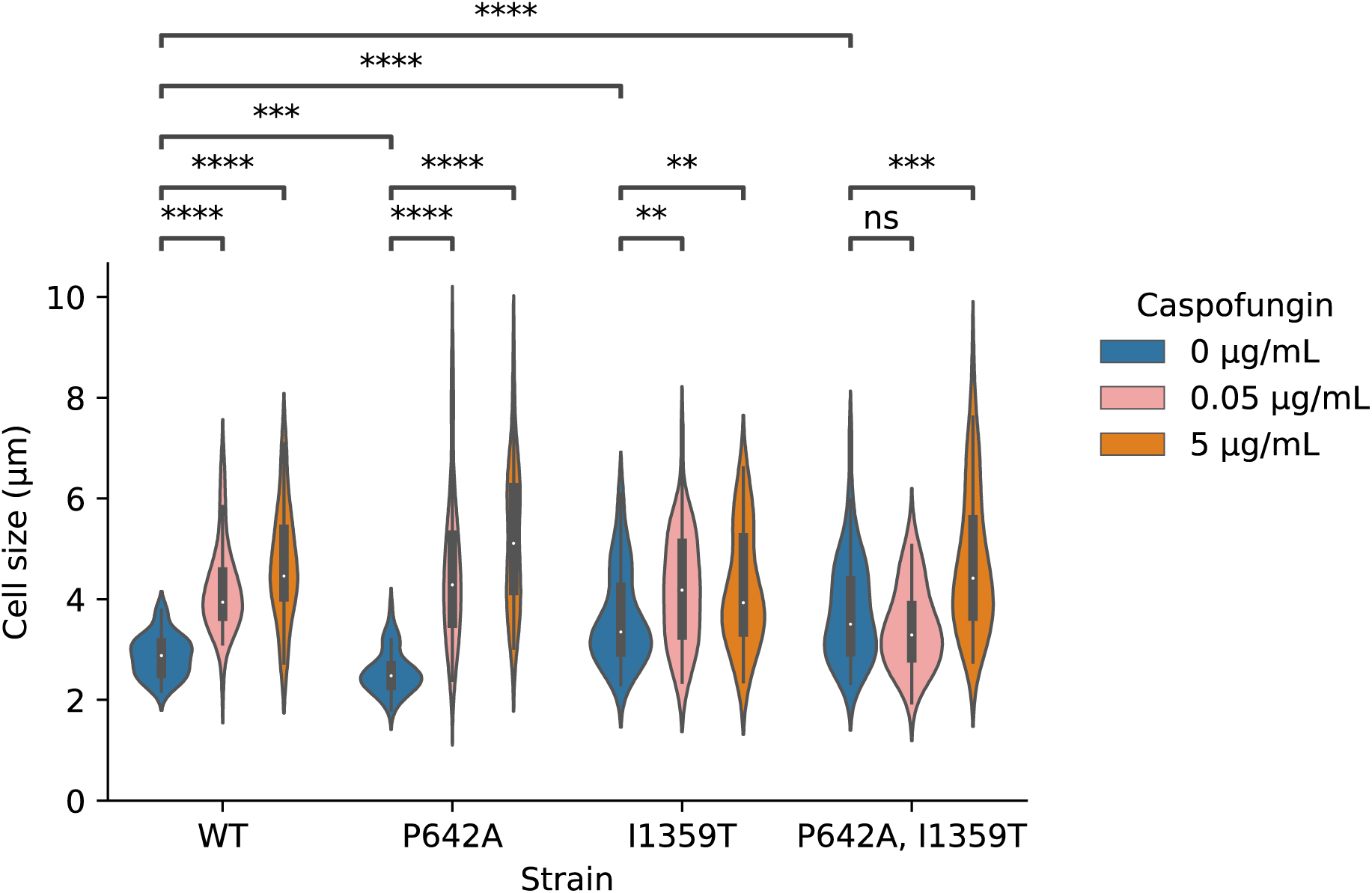
Violin-plot of the cell sizes of *C. lusitaniae* strains. The distribution of cell sizes (short axis) was measured for WT, P642A, 11359T and P642A, 11359T cells without caspofungin and after caspofungin treatment at 0.05 µg/mL and 5 µg/mL (n = 5O cells from 3 independent experiments; Box percentile: 25^th^-75^th^; White dots correspond to median).

### Susceptibility to other cell wall and osmotic stresses

Morphological changes under caspofungin treatment, particularly cell swelling, led us to test their susceptibilities to other compounds that interfere with the fungal cell wall and to induce rescue pathways. We used calcofluor white (CFW) and Congo red (CR), whose damage to the cell wall activates the CWI pathway; caffeine, which activates the CWI pathway without damaging the cell wall; and sorbitol and NaCl, to test the HOG response (Fig. 6). WT and mutant strains had comparable susceptibilities to the hyperosmotic stress generated by sorbitol (MIC 1500 mM) and NaCl (MIC 1250 mM), and to the stress caused by caffeine (MIC 25 mM). In contrast, the mutant strains exhibited enhanced susceptibilities to CFW and CR relative to the WT. The P642A substitution in Fks1p increased the susceptibility of the strain four-fold to CFW and eight-fold to CR. The I1359T substitution, with or without P642A, increased susceptibility eight-fold for CFW and 32-fold for CR (Fig. 6). The different responses of the Fks1p mutants to cell wall stressors (caspofungin, CFW, and CR) prompted us to explore the main cell integrity pathways possibly related to caspofungin-induced cell wall stress in the Fks1p mutants, *i.e.*, the calcineurin, HOG, and CWI pathways.

**Figure 6.**
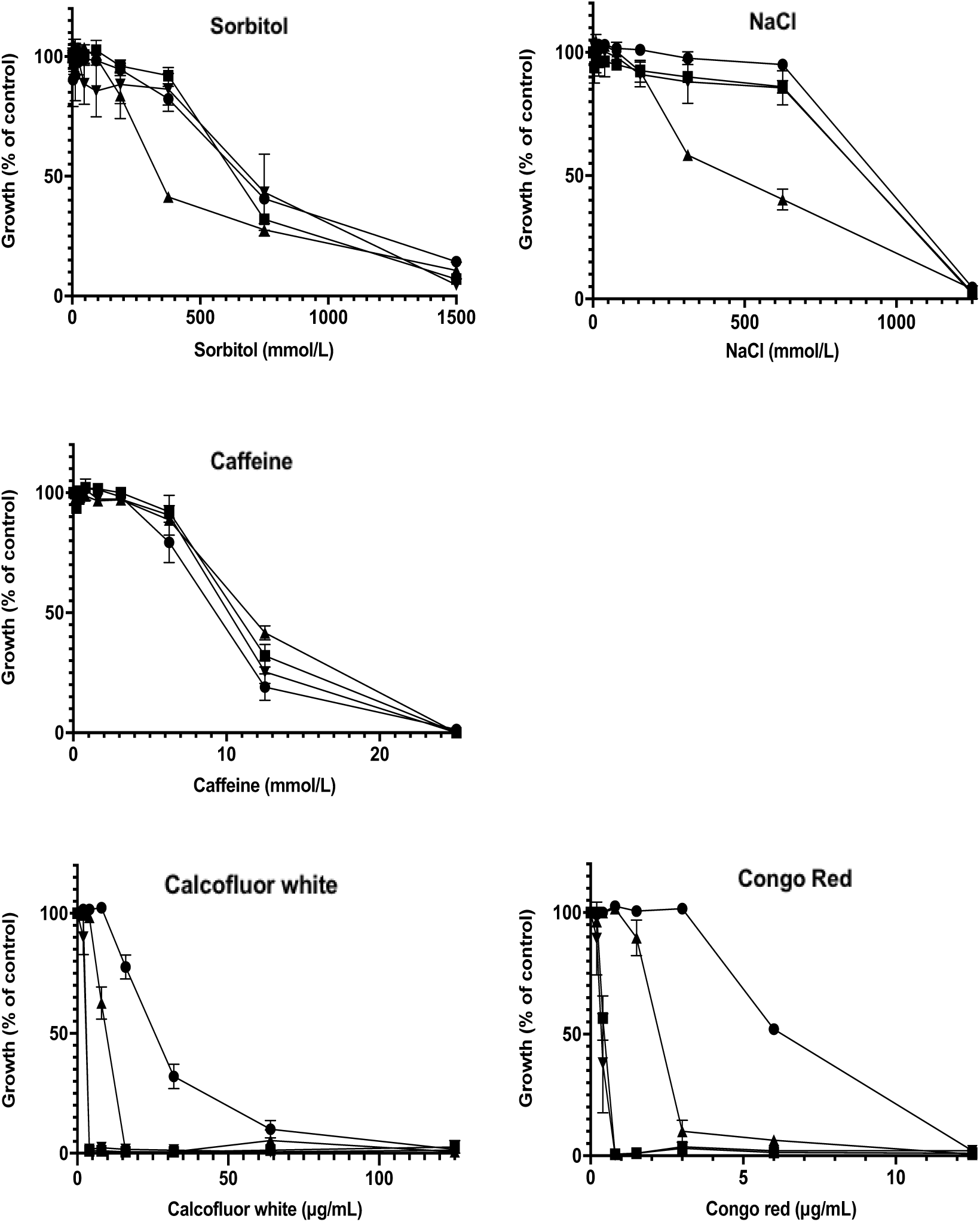
Susceptibility of *C. lusitaniae* WT and mutant strains to osmotic stress and cellular and cell wall stressors. M1C was considered as the concentration of the stressor which inhibited 8O to 1OO% of the growth of the strains compared to their growth in the culture medium without stressor after 48 h of incubation. WT (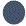); P642A (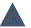); 11359T (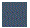); P642A, 11359T (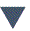).

### Transcriptional analysis of target genes of cell wall synthesis and integrity in *C. lusitaniae*

The expression levels of the genes encoding chitin synthase (*CHS1, CHS2, CHS3, CHS8*), glucan synthase (*FKS1*), the catalytic subunit of calcineurin (*CNA1*), and the major mitogen-activated protein kinases of the HOG (*HOG1*) and CWI (*MKC1*) pathways, were analyzed by quantitative RT-PCR in WT and mutant strains cultivated in drug-free RPMI medium and in caspofungin-supplemented medium (5 µg/mL for 3 h). The expression level of each gene in the WT strain in drug-free medium was normalized to 1 by reference to *ACT1* mRNA. The expression levels of the genes according to strain and growth conditions were compared to the WT in drug-free medium; the variations did not exceed three-fold. Only significant changes (p ≤ 0.05) in the means of three independent biological replicates are discussed below (Fig. 7). Caspofungin treatment of WT *C. lusitaniae* increased the expression levels of *CHS1, CHS2*, *CHS3*, *CNA1*, and *FKS1*. The P642A substitution in Fks1p resulted in enhanced expression of *CHS3*, *CHS8*, *CNA1*, *FKS1* and *HOG1* in drug-free medium; the addition of caspofungin further increased the expression level of *CHS3*, but that of *FKS1* was unaffected. The I1359T substitution alone, and in combination with P642A, resulted in enhanced expression of *CHS3*, *CHS8*, *CNA1*, and *MKC1* in drug-free medium. Adding caspofungin increased the expression of *CHS1*, *CHS2*, and *CHS3* in the I1359T mutant, and of *CHS2* in the double mutant expressing the P642A and I1359T substitutions. To summarize, the transcriptional response to caspofungin involved increased expression of the calcineurin, chitin synthase, and glucan synthase genes, except in strains with the I1359T substitution, in which *FKS1* was not over-expressed to the benefit of *MKC1*. Accordingly, when compared to WT strain, I1359T tends to decrease the expression of *FKS1* even in the presence of caspofungin and leads to increase the expression of *MKC1*, even in the absence of caspofungin.

**Figure 7.**
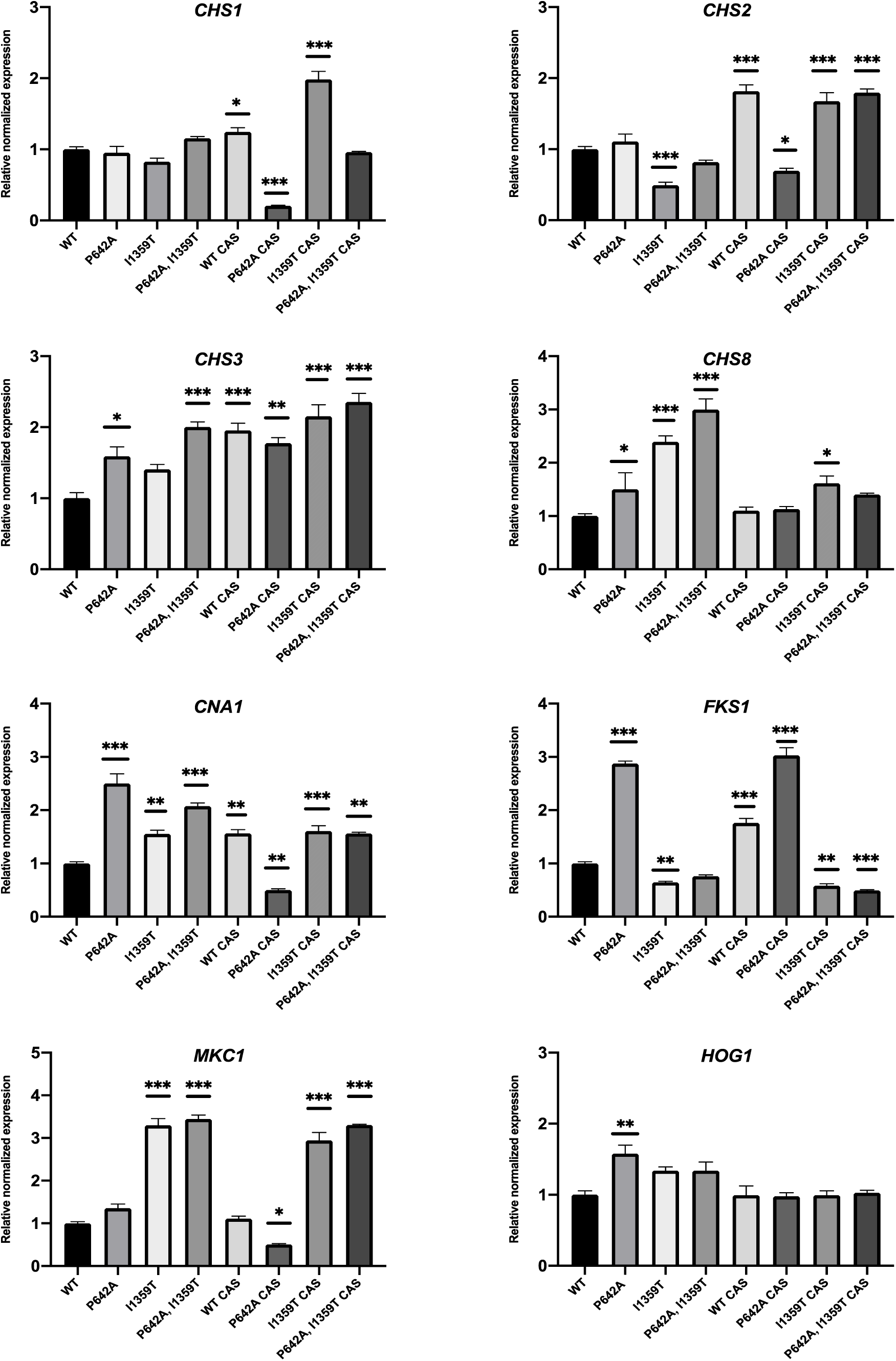
Quantitative transcriptional analysis of genes involved in cell wall biosynthesis and in cellular rescue pathways in *C. lusitaniae* WT and mutant strains exposed or not to caspofungin (CAS, 5 µg/mL, 1 h). * p<0.05; ** p<0.01; *** p<0.0O5.

### Mkc1p is activated in strains expressing I1359T Fks1p even in the absence of caspofungin

Monoclonal antibodies against phosphorylated Mkc1p, phosphorylated Hog1p, and tubulin were used for western blot analysis of protein extracts of the *C. lusitaniae* WT and mutant strains cultivated in the absence and presence of caspofungin (Fig. 8). For the HOG pathway, strains expressing Fks1p with the I1359T substitution exhibited an increased content of phosphorylated Hog1p under caspofungin exposure. Compared with the WT, a very small quantity of phosphorylated Hog1p was also detected in strains with Fks1p polymorphisms cultivated in drug-free medium. For the CWI pathway, as expected from previous findings (16), all of the strains analyzed responded to caspofungin by phosphorylating Mkc1p. Interestingly, phosphorylated Mkc1p was detected in protein extracts from the I1359T Fks1p mutants, with or without P642A, cultivated in drug-free medium. Constitutive activation of the CWI pathway strongly suggests that the I1359T substitution is deleterious to the cell.

**Figure 8.**
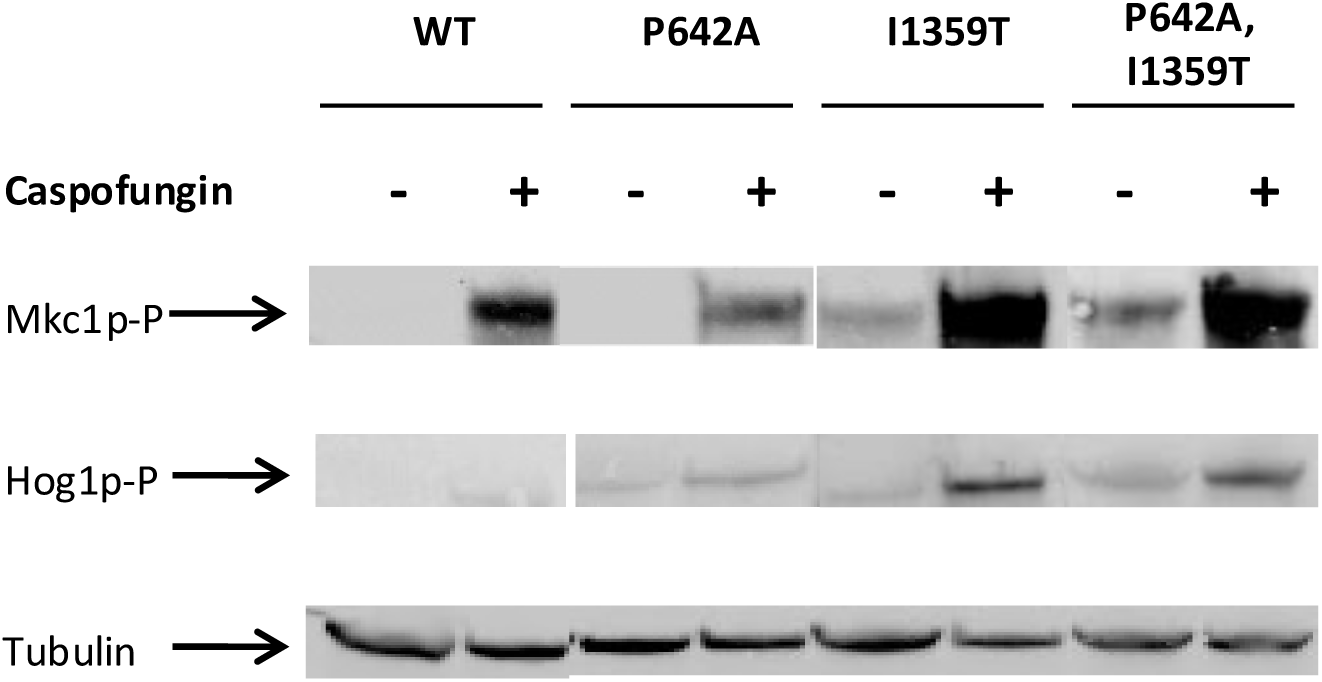
Western blot analysis of the phosphorylated MAP kinases Mkc1p-P and Hog1p-P in crude protein extracts of *C. lusitaniae* WT and *FKS1* mutant strains exposed or not to caspofungin (5 µg/mL, 1 h). All the samples were analysed on the same blot.

### Deletion of *MKC1* in *C. lusitaniae* strains expressing Fks1p with the I1359T substitution restores caspofungin susceptibility *in vitro*

The ORF OVF10720.1 of strain CBS 6936 of *C. lusitaniae* (29) encodes a putative Mkc1p of 546 amino acids having 79.8% identity with the *C. albicans* ortholog protein CR_00120C_A (30), and having the conserved TEY phosphorylation motif at amino acids 193-195. *MKC1* loss-of-function mutants were obtained in the WT and mutant *FKS1* strains of *C. lusitaniae* expressing the I1359T substitution with or without P642A by targeted integration of an *URA3* deletion cassette at the *MKC1* locus. Strains with a reconstituted *MKC1* locus were also generated for each null mutant. The strains were tested for caspofungin susceptibility (Table 1). Deletion of *MKC1* in the WT had little effect on caspofungin susceptibility, decreasing the MIC from 0.125 to 0.0625 µg/mL. In contrast, deletion of *MKC1* in the strain expressing Fks1p with the I1359T substitution restored caspofungin susceptibility to the WT level (0.125 µg/mL), and to the level of the P642A mutant in the strain expressing Fks1p with both the P642A and I1359T substitutions (0.75 µg/mL). In each case, the phenotype observed was reversed by reintegrating a WT *MKC1* allele in each null mutant. This finding demonstrates that caspofungin resistance mediated by the I1359T substitution is mainly dependent on *MKC1* and on activation of the CWI rescue pathway *in vitro*.

**Table 1.**
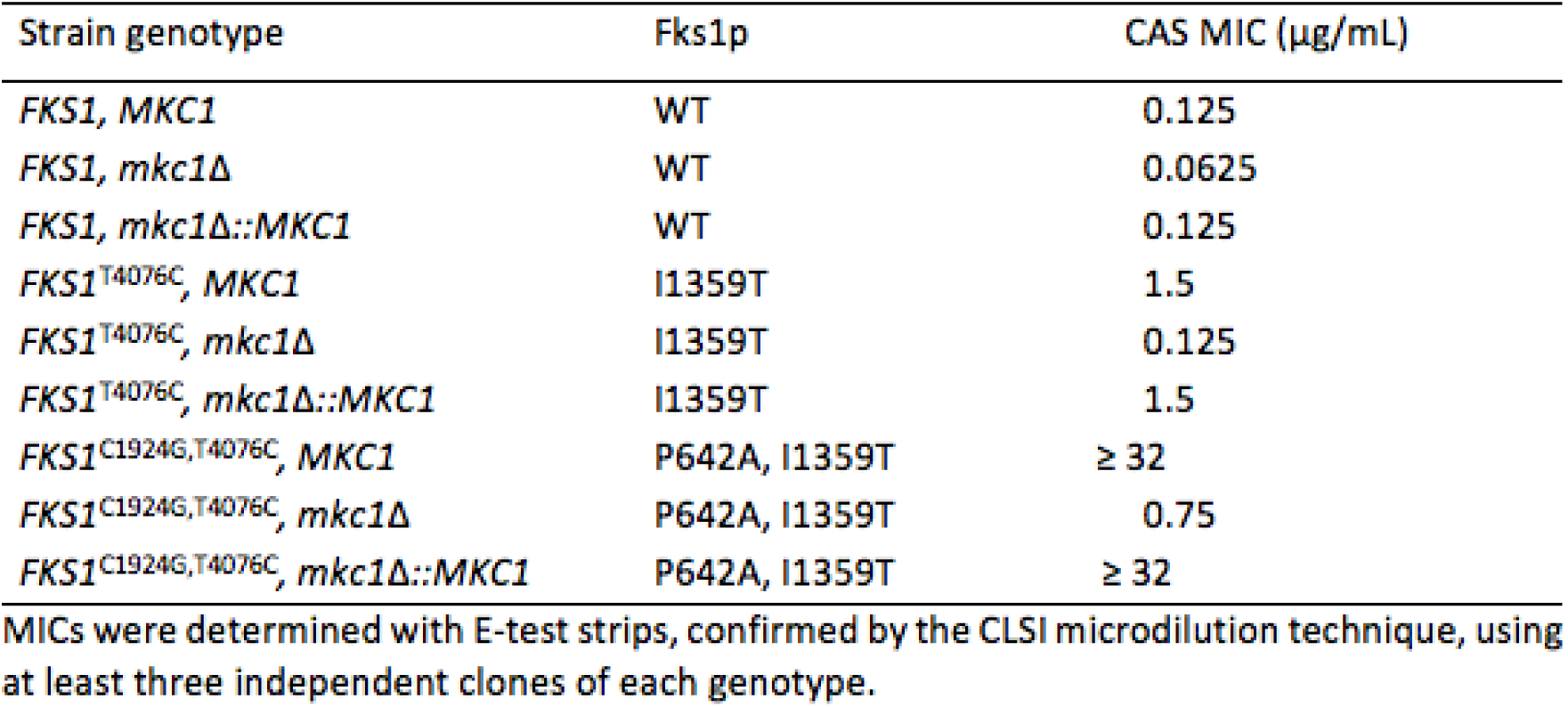
Caspofungin MICfor 1sogenle WT and mutant:*fKS.1* strains of C. *fusitaniae* in wild type, deleted and re-constructed genetic badkgrounds for *MKCl*.

| Strain genotype | Fks1p | CAS MIC (µg/mL) |
| --- | --- | --- |
| *FKS1*, *MKC1* | WT | 0.125 |
| *FKS1*, *mkc1*Δ | WT | 0.0625 |
| *FKS1*, *mkc1*Δ::*MKC1* | WT | 0.125 |
| *FKS1*<sup>T4076C</sup>, *MKC1* | I1359T | 1.5 |
| *FKS1*<sup>T4076C</sup>, *mkc1*Δ | I1359T | 0.125 |
| *FKS1*<sup>T4076C</sup>, *mkc1*Δ::*MKC1* | I1359T | 1.5 |
| *FKS1*<sup>C1924G,T4076C</sup>, *MKC1* | P642A, I1359T | ≥ 32 |
| *FKS1*<sup>C1924G,T4076C</sup>, *mkc1*Δ | P642A, I1359T | 0.75 |
| *FKS1*<sup>C1924G,T4076C</sup>, *mkc1*Δ::*MKC1* | P642A, I1359T | ≥ 32 |

MICs were determined with E-test strips, confirmed by the CLSI microdilution technique, using at least three independent clones of each genotype.

### Effects of *FKS1* allelic variability on the fitness of *C. lusitaniae in vitro*

The effects of the *FKS1* mutations on fitness were evaluated using an *in vitro* model of *Candida*-macrophage interactions (31). J774 murine macrophages were infected at a multiplicity of infection (MOI) of 1: 1 with *C. lusitaniae* WT and *FKS1* mutants, and cellular interactions were measured at 30 min, 5 h, and 24 h post-infection. Prior to infection, yeasts were treated or not with caspofungin at low (0.05 µg/mL) and high (5 µg/mL) concentrations.

The first parameter analyzed was the fraction of macrophages engaged in phagocytosis. Without caspofungin, the proportion of macrophages engaged in phagocytosis was 44 to 66% for all strains at 5 h and 24 h post-infection (Fig. 9 at 24 h). Treatment of WT cells with caspofungin at 0.05 µg/mL reduced by 50% the number of macrophages involved in phagocytosis at each time point. The percentage of macrophages phagocytosing the single I1359T and double P642A, I1359T mutants also decreased by half after treatment with caspofungin at 5 µg/mL (Fig. 9).

**Figure 9:**
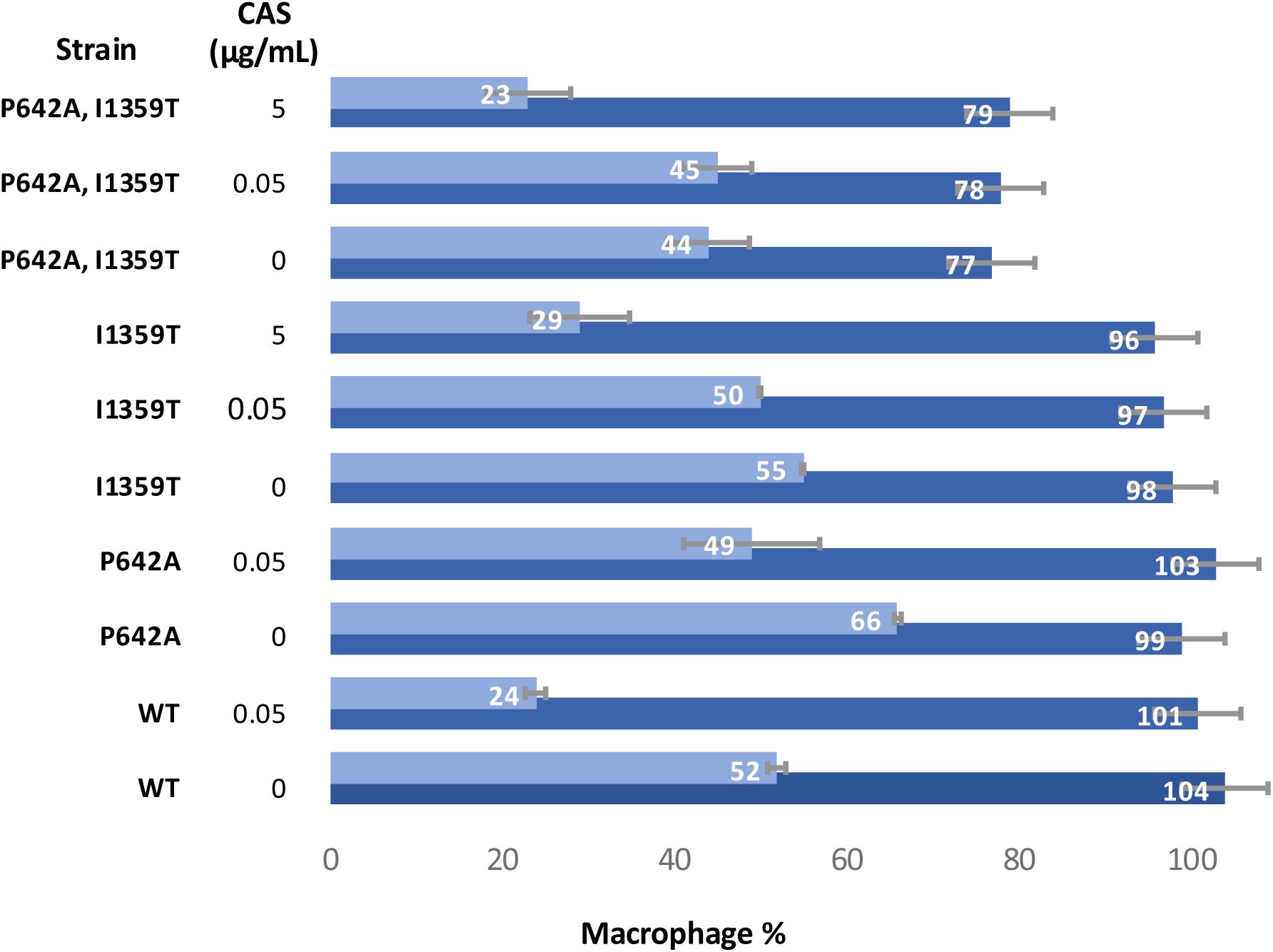
Macrophage phagocytosis assays of WT and *FKSl* mutants of *C. lusitaniae*. Yeasts exposed or not to different concentrations of caspofungin (CAS) and J774 murine macrophages were mixed at a MO1 of 1Y:1M and the percentage of living macrophages (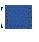) and of phagocytosing macrophages (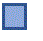) was measured 24 h post-infection by flow cytometry (± SEM from 3 independent experiments performed in quintuplets).

The second parameter was the fungal biomass internalized by the macrophages. Without caspofungin treatment, the fungal biomass phagocytozed at 24 h post-infection was increased significantly (∼ 10-fold) for the two mutants expressing the I1359T Fks1p, compared to those expressing WT and P642A Fks1p (Fig. 10). Exposure to caspofungin increased the fungal biomass ingested by phagocytes, noticeably for *C. lusitaniae* WT cells exposed to 0.05 µg/mL caspofungin and for the single I1359T and double P642A, I1359T mutant cells exposed to 5 µg/mL caspofungin. Although the phagocytosed fungal biomass may have been overestimated in the mutants due to their larger size and changes in cell wall content, these results suggest that the I1359T Fks1 protein promotes recognition of yeasts by macrophages and improves phagocytosis efficiency under antifungal pressure, because fewer phagocytes (Fig. 9) were recruited but more yeasts were engulfed (Fig. 10).

**Figure 10.**
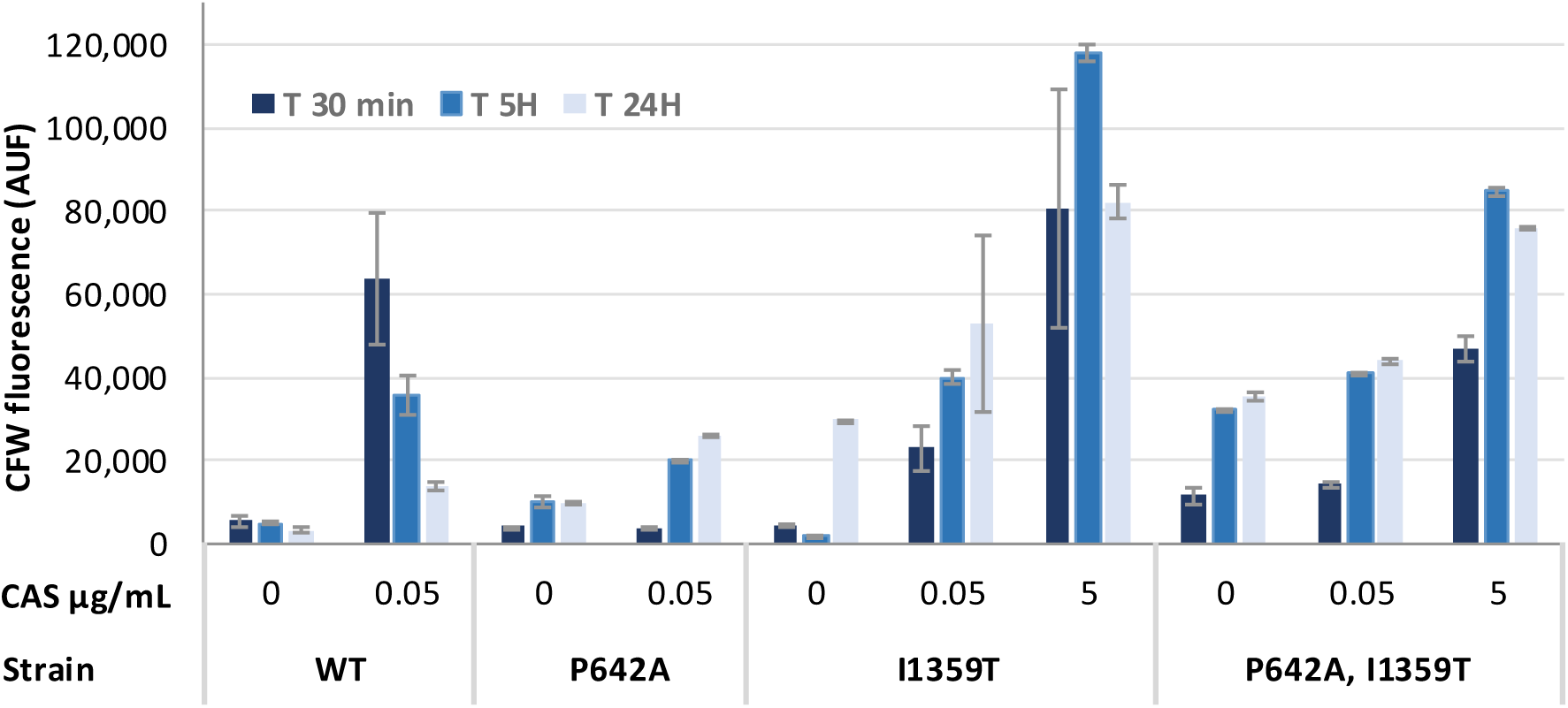
Relative mass of WT and *FKS1* mutants of *C. lusitaniae* phagocytosed by murine J774 macrophages. CFW-labeled yeasts exposed or not to different concentrations of caspofungin (CAS) and J774 murine macrophages were mixed at a MO1 of 1Y:1M. Mean fluorescence intensity (MF1) of intra-macrophagic yeasts after quenching with trypan blue was measured 24 h post-infection (± SEM from 3 independent experiments performed in quintuplets). AUF: arbitrary units of fluorescence.

The third parameter was the survival of intra-macrophagic yeasts at 24 h post-infection (Fig. 11). Interestingly, under drug-free conditions, survival rates were higher for yeasts expressing Fks1 protein with the I1359T substitution, with or without P642A, compared to WT and P642A. At 0.05 µg/mL caspofungin, survival was higher for I1359T relative to the WT, and was comparable to the P642A Fks1p strain.

**Figure 11.**
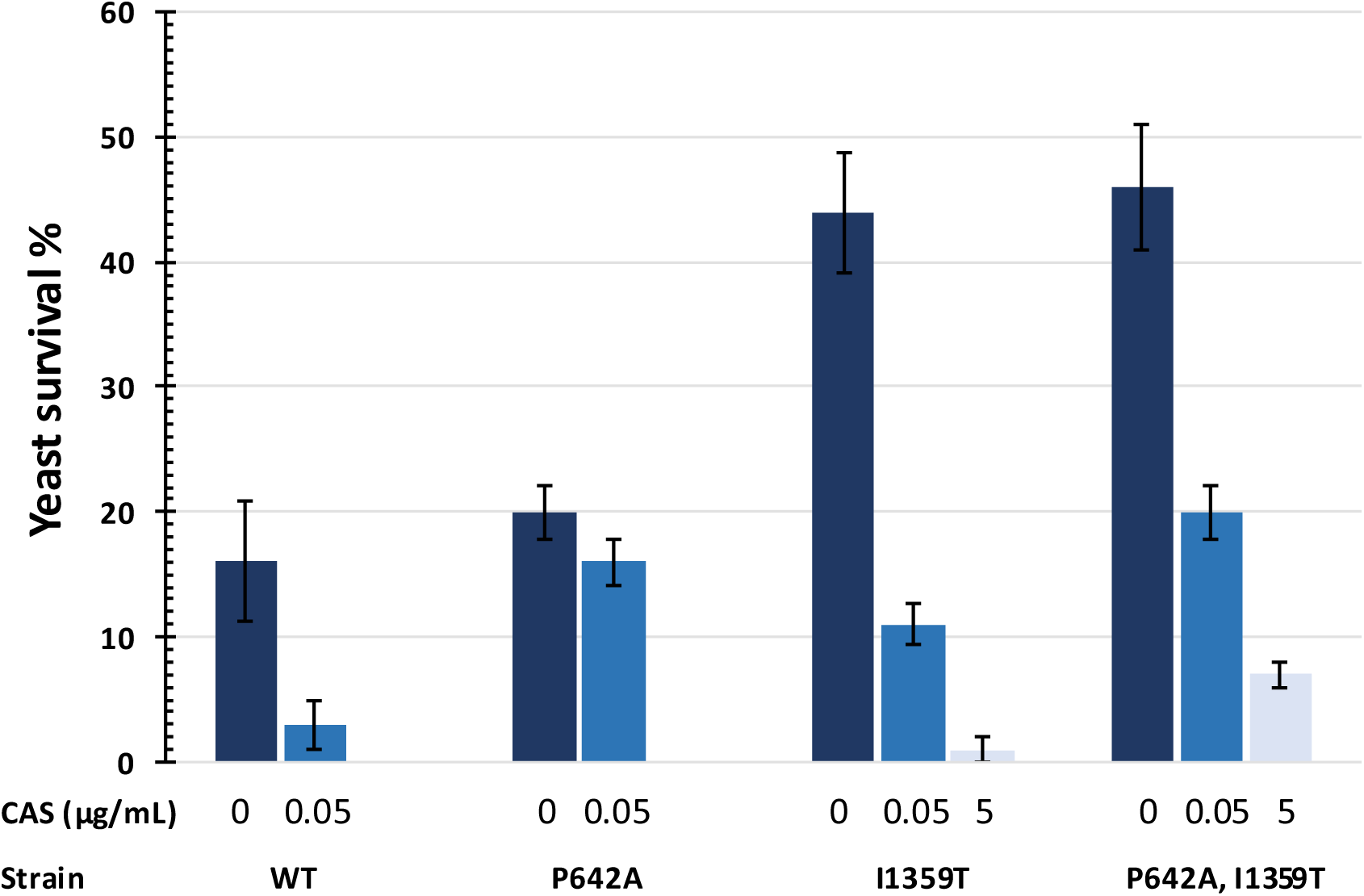
1ntramacrophagic survival of WT and *FKSl* mutants of *C. lusitaniae*. Yeasts exposed or not to different concentrations of caspofungin (CAS) and J774 murine macrophages were mixed at a MO1 of 1Y:1M and incubated 24 h. After macrophage lysis, survival percentage was obtained by the number of CFU after plating 1OO yeasts on YPD (± SEM from 3 independent experiments).

## DISCUSSION

After collecting a clinical blood isolate of *C. parapsilosis* resistant *in vitro* to echinocandins having a new mutation in *FKS1*, we characterized the cellular and molecular effects of this mutation in *C. lusitaniae*. The *C. parapsilosis* isolate carried a mutation in one of the two *FKS1* alleles, resulting in the replacement of an isoleucine by a threonine at position 1380 in β-1,3-glucan synthase. This polymorphism was recently reported in an *FKS1* allele of *C. albicans* resistant to both anidulafungin and ibrexafungerp (32). In this work, we provide evidence that it contributes to echinocandin resistance *in vitro* after its expression in the haploid yeast *C. lusitaniae* (10). We chose to study the effect of the mutation in a genetic context different from the one in which it was initially isolated, in order to eliminate any genetic and/or epigenetic factors that might have contributed to its selection. The presence of the P660A polymorphism in *C. parapsilosis* is a genetic factor that contributed to the selection of the I1380T mutation. By expressing I1380T in the presence and absence of P660A in the *C. lusitaniae* model, we were able to demonstrate that it is indeed the combination of these two polymorphisms that confers a high level of *in vitro* resistance to echinocandins, rather than the I1380T mutation alone. In *C. albicans*, most of the resistance mutations are concentrated in two conserved regions of the *FKS1* gene, the so-called hot spot 1 (HS1) and 2 (HS2) regions (12, 33). These regions presumably bind the cyclic peptidic part of echinocandins (34). A third conserved region described in *S. cerevisiae*, HS3, located between HS1 and HS2, could be involved in the binding of the hydrophobic acyl tail of echinocandins (34). In the *C. parapsilosis* group, other than the naturally occurring P660A polymorphism and the recently described mutations R658G (14) and F652S (13) in HS1, most of the mutations suspected to be involved in echinocandin resistance were located outside the HS1 and HS2 regions (15). The I1380T mutation described here is located four amino acids downstream of HS2 in a peptidic region expected to be exposed outside the plasma membrane (34). Replacing the nonpolar isoleucine with the similar-in-size but polar threonine can have marked consequences. Threonine has a hydroxyl group that can be involved in numerous reactions, such as phosphorylation, *O*-linked glycosylation, proteolytic cleavage, and hydrogen binding to other amino acids, which can influence the secondary and tertiary structures of proteins, and has been implicated in protein-protein interactions (35) (36).

It is possible that the I1380T amino acid substitution reduces the binding affinity of the HS2 domain to echinocandins. However, when expressed in *C. lusitaniae* at the equivalent position, and compared to the WT Fks1 protein, the I-T substitution alone conferred a 12-fold increase in the caspofungin MIC (from 0.125 to 1.5 µg/mL), and a 37-fold increase in the caspofungin IC_50_ value. The I1380T substitution must be associated with the P660A polymorphism of HS1, which itself has a moderate effect on caspofungin MIC, to confer a high caspofungin MIC (≥ 32 µg/mL, comparable to the clinical CPAR isolate) and a 270-fold increase in the caspofungin IC_50_ value compared to the WT. These findings indicate that both the HS1 and HS2 domains of Fks1p contribute to binding caspofungin, and therefore that they are in conformational close vicinity. The P-A and I T substitutions, in combination, conferred the same level of pan-echinocandin resistance *in vitro* in both *C. parapsilosis* and *C. lusitaniae*. Therefore, the conformational changes must involve conserved regions in Fks1p between the two species. A cryo-electron microscopy study of Fks1p in *S. cerevisiae* (37) revealed that HS1 and HS2 are located in two neighboring transmembrane helices, TM5 for HS1 and TM8 for HS2, on the outer side of the cell membrane. Also, P642 and I1359 (*C. lusitaniae* Fks1p coordinates) are on peptide segments close enough to interact. The authors suggested that any substitutions at or near these positions may contribute to a substantial change in the lipid environment of the enzyme, sufficient to reduce the binding affinity to caspofungin. Moreover, TM8 amino acids contribute to the β-glucan translocation channel and some interact with the newly synthesized glucan chain. In *S. cerevisiae* Fks1p, the aromatic F1363, just upstream of I1364 and corresponding to I1359 in *C. lusitaniae*, is expected to bind glucans (37).

Decreasing the binding capacity of Fks1p to caspofungin is not the only effect of the combination of P660A and I1380T mutations. Using vibrational spectroscopy, we observed that the *C. lusitaniae* strains expressing I1359T Fks1p (equivalent position of I1380T), especially with P642A (equivalent position of P660A), had the lowest 1,3-ß-glucan contents in the cell wall compared to the WT when exposed to high caspofungin concentration. Although caspofungin has a decreased binding capacity to I1359T Fks1p, it appears to exhibit increased inhibitory activity, as less glucan is synthesized by I1359T Fks1p.

In *C. albicans* and *C. glabrata*, some amino acid substitutions contributing to caspofungin resistance concomitantly decreased the *Vmax* of Fks1p (33, 38, 39). Unlike a report that the glucan content in the cell wall of echinocandin-resistant *C. albicans* strains was unaffected by Fks1p mutations (38), our work shows a direct correlation between the Fks1 I1359T polymorphism and the glucan deficit in the cell wall of *C. lusitaniae*.

Further evidence of abnormal Fks1p activity was provided by observation of the cell surface by AFM. Some of the yeasts expressing I1359T Fks1p, with or without P642A, had irregular rough surfaces, and were larger than the WT, in drug-free medium. This is a phenotype similar to WT cells exposed to low concentrations of caspofungin. This phenotype can be the consequence of a defect of glucan synthase activity, mainly caused by I1359T, because the P642A substitution has no effect on cell wall phenotype. Our findings confirm previous reports that yeast cell integrity in the presence of echinocandins may in part be ensured by increased chitin synthesis (16, 17, 19, 20, 38, 40). Cells were larger and had a reduced cell wall glucan content but did not exhibit greater susceptibility to osmotic stress. The I1359T mutants had no particular sensitivity to caffeine. This purine alkaloid is capable of activating the Pkc1p-Mkc1p CWI pathway (30) via TOR complex 1, but does not affect cell wall remodeling because of its inability to activate the transcription factor Rlm1 (41).

Lack of susceptibility to caffeine suggests that the CWI pathway was active in the I1359T mutants. However, these mutants exhibited a marked growth defect in the presence of CFW and CR. Both chemicals have similar effects, which can partially mimic those of echinocandins, by binding and inhibiting the synthesis of glucans and chitin at the cell wall (41, 42). Their damage is perceived by cell wall sensors, which activate the CWI pathway up to the transcription factor Rlm1, thereby upregulating the expression of the genes encoding glucan and chitin synthases. The greater susceptibility of the I1359T mutants to these inhibitors indicated that at least one component of the cellular response to cellular damage was dysfunctional. Targeted transcriptional analysis in *C. lusitaniae* of the main genes involved in signaling and cell wall synthesis confirmed, as expected from previous reports, the enhanced expression of calcineurin A (16), chitin synthase, and *FKS1* (43) in cells exposed to caspofungin. Interestingly, increased expression of *FKS1* was not observed in strains with the I1359T Fks1p mutation, in which *MKC1* was upregulated. Implication of this MAP kinase was confirmed at the protein level: the phosphorylated active form of Mkc1p was markedly increased in all the strains exposed to caspofungin, and in the I1359T mutants cultivated in drug-free medium. Restoration of caspofungin sensitivity following deletion of *MKC1* in *C. lusitaniae* mutants with I1359T Fks1p revealed that resistance *in vitro* mediated by I1359T Fks1p is dependent on *MKC1* and on activation of the CWI pathway. The direct involvment of *MKC1* in echinocandin resistance could be demonstrated in this study by using the haploid *C. lusitaniae* yeast model, as, unlike the *C. parapsilosis* clinical isolate, there is no wild-type allele to compensate for the defect in the I1359T Fks1 glucan synthase. Although the I1359T mutant cells were larger in drug-free medium, the weak Hog1p phosphorylation, which is involved in adaptation of the cell to high-osmolarity conditions, was not specific to I1359T, as it was also observed in P642A. Activation of the CWI pathway without exposure to echinocandin likely results from defective I1359T Fks1p activity, which is concordant with the damage seen by AFM on the cell wall of the mutant even in drug-free medium, and with the lowest glucan content detected by IR spectroscopy. This defect may also explain the greater susceptibility to CFW and CR in I1359T Fks1p mutants, which cannot compensate for the damage to glucans by upregulating the expression of a fully functional Fks1 enzyme. It may also explain the apparent loss of fitness in *C. lusitaniae* strains expressing I1359T Fks1p during interactions with J774 macrophages. Compared with the WT under caspofungin treatment, 50% fewer phagocytes were recruited to engulf a significantly greater fungal biomass of yeasts expressing I1359T Fks1p. The significant decreases in the ß-glucan contents of the mutants could enhance the exposure of mannose and glycoproteins, which are recognized by macrophage TLR2, TLR4, and C-type Lectins receptors (44), and/or to unmasking of chitin, the proportion of which in the cell wall is increased by antifungals. Chitin reinforcement in the cell wall may explain the enhanced intra-macrophagic survival of I1359T Fks1p mutants, either *via* the NLR cytoplasmic receptor NOD2 and its interaction with chitin, mannose receptor, and TLR9, leading to the production of IL-10 (44); or *via* chitin-mediated inhibition of nitric oxide synthase (45). Loss of fitness was reported in resistant *C. albicans* strains, in which several Fks1p mutations affected growth rate, filamentation, and virulence in a variety of animal models (38).

In conclusion, when expressed in *C. lusitaniae*, the mutation P642A, which corresponds to the P660A species-specific polymorphism of *C. parapsilosis*, had little effect on the caspofungin MIC, and no effect on glucan content of the cell-wall or survival after macrophage phagocytosis. These results support clinical practice, in which echinocandins at standard concentrations are used for the treatment of most *C. parapsilosis* invasive infections (12). In contrast, I1359T is a deleterious mutation that decreases the binding capacity of caspofungin to Fks1p and also impairs glucan synthesis. The resulting yeast cells have abnormal cell walls, and exhibit constitutive activation of the CWI rescue signaling pathway, notably by the phosphorylation of Mkc1p. We have finally demonstrated that Mkc1 is the main molecular determinant of caspofungin resistance *in vitro* in *C. lusitaniae* yeasts expressing Fks1p I1359T. However, we believe that the activation of rescue signaling pathways to resist to echinocandin is likely not confined to the Fks1p I1359T mutation. Recently, we showed that rapamycin restored caspofungin susceptibility *in vitro* in resistant strains of *C. albicans* with the Fks1p S645P mutation, as well as in *C. lusitaniae* strains expressing the equivalent mutation (46). Notably, rapamycin activates Mpk1p, the ortholog of Mkc1p in *S. cerevisiae* (47), suggesting that other factors in the signaling network, beyond Mkc1p, may be involved in echinocandin resistance, depending on the mutation affecting Fks1p.

## MATERIALS AND METHODS

### *Candida* strains and culture conditions

The *C. parapsilosis* clinical strain CPAR was identified by matrix-assisted laser desorption ionization– time of flight mass spectrometry Microflex LT systems (Bruker Daltonics), with FlexControl (version 3.0) software (Bruker Daltonics). All of the *C. lusitaniae* strains constructed in this study or from previous works (10) (48) were derived from the WT reference strain CBS 6936 (ATCC 38553). All of the strains and their genotypes are listed in Supplemental Data S5. Yeasts were routinely cultivated in YPD medium and on YNBS after transformation by electroporation, as described previously (10). RPMI 1640 medium (Sigma-Aldrich) supplemented with 2% glucose and buffered to pH 7.0 with 0.165 M MOPS was used to test susceptibility to antifungals and other stressor molecules.

### Nucleic acid extraction and PCR amplification

Yeast DNA was extracted using a glass-bead method (48) and PCR was performed using the high-fidelity DNA polymerase Pfu (Promega). RNAs were extracted from 1 × 10^8^ cells suspended in 1 mL of TRI REAGENT® (Molecular Research Center) and disrupted using the glass-bead method (49). RNA quality and quantity were evaluated on an Agilent Bioanalyzer with RNA 6000 Nano Chips. For RT-qPCR, 50 ng of RNA from each sample were amplified using the GoTaq One-Step RT-qPCR System (Promega) with the SYBR-Green intercalating agent on the Bio-Rad CFX Thermocycler. Relative quantification of gene expression was carried out by the 2^-ΔΔCt^ method (50) using *ACT1* as the reference gene, in triplicate experiments using independent RNA samples. Fluorescence data from RT-qPCR were collected and analyzed, including statistical tests, with CFX Manager software, v. 3.1 (Bio-Rad). The sequences of the primers used for PCR and RT-qPCR and the GenBank accession numbers of the target genes are provided in Supplemental Data S6.

### Construction of *C. lusitaniae FKS1* mutants

*FKS1* allele replacement was achieved in *C. lusitaniae* as described previously (10), by introducing into strain F1 *trp1*Δ3’, *ura3*Δ5’ two DNA fragments covering the entire *FKS1* ORF. Both fragments overlap a 40 bp region where SNPs of interest were inserted. In this study, the single mutant CLUS I1359T and the double-mutant CLUS P642A, I1359T were constructed starting from the genomic DNA of strains F1 *TRP1, URA3*, and CLUS Fks1p P642A, respectively, as described previously (10). The following mutations were introduced separately and in combination in the *FKS1* gene of *C. lusitaniae*: C1924G (codon change CCT to GCT) for P642A and T4076C (codon change ATT to ACT) for I1359T, corresponding to the coordinates C1978G (P660A) and T4139C (I1380T) in the *FKS1* gene and protein of *C. parapsilosis*. *FKS1* nucleotidic changes were verified by DNA sequencing (Eurofins Genomics). The primers are listed in Supplemental Data S6.

### Construction of *C. lusitaniae mkc1* deletion mutants

The *mkc1* deletion mutants were generated using a DNA cassette of *URA3* flanked by the 5’– and 3’-UTR regions of *MKC1*. The recipient strains for transformation were 6936 *ura3*Δ expressing a WT *FKS1* allele, and strains 1359T *ura3*Δ and P642A/I1359T *ura3*Δ expressing mutant *FKS1* alleles. Reconstructed strains were obtained after transformation with a PCR DNA fragment containing a *MKC1* allele, and selection on YNBS supplemented with 5-fluoroorotic acid (0.8 mg/mL) and uracil (50 µg/mL). Deletion/insertion of *MKC1* was verified using PCR and nucleotide sequencing (Eurofins Genomics). The primers are listed in Supplemental Data S6.

### Susceptibility testing

Susceptibility to antifungals was determined by Etest (bioMérieux) according to the manufacturer’s instructions. MICs were read after incubation for 48 h at 35°C. For caspofungin supplementation, we used caspofungin acetate powder (Merck Sharp and Dohme; Kenilworth, USA). Susceptibility to toxic compounds was evaluated using microdilution assays adapted from CLSI standards (51) in RPMI 1640 medium. Yeast growth was measured after incubation for 48 h at 35°C using a microplate reader at 450 nm. Experiments were conducted at least in triplicate. Calcofluor white (CFW), Congo red (CR), and caffeine were purchased from Sigma-Aldrich.

### Inhibition measurement of glucan synthase

Yeast cells were grown to early stationary phase in YPD medium and collected by centrifugation. Cell disruption, membrane protein extraction, and partial 1,3-β-glucan synthase purification by product-entrapment were performed as described previously (39). Reactions were initiated by the addition of product-entrapped glucan synthase. Sensitivity to caspofungin was measured by polymerization assay using a 96-well 0.65 µm multiscreen HTS filtration system (Millipore Corp.) in a final volume of 100 µL, as described previously (11). Serial dilutions of the drugs (0.01 to 10,000 ng/mL) were used as calibration standards. Inhibition profiles and IC_50_ values were determined using a normalized response (variable-slope) curve fitting algorithm with Prism v. 8.1.2 software (GraphPad Software).

### Atomic force microscopy imaging

AFM experiments were performed at room temperature (20°C) using a BioScope Resolve AFM (Bruker) and MSCT tips (MicroLever, Bruker). Cells were cultivated in RPMI medium with (0.05 or 5 µg/mL) or without caspofungin for 16 h at 30°C with agitation. Next, 200 µL of each suspension were deposited for 2 h on glass substrates coated with a thin layer of Cr (∼10 nm) and Au (∼30 nm).

After sedimentation, surfaces were gently rinsed in ultrapure water and dried overnight at 30°C. Images were taken in air to enhance the morphological differences induced by the drug. Cell size was determined by measuring the short axis of 50 cells selected randomly in images from three independent experiments for each condition.

### IR-ATR

IR-ATR spectra were recorded between 4000 and 800 cm^-1^ on a Bruker Vertex 70v spectrometer as described previously (17). The resolution of the single beam spectra was 4 cm^-1^. A nine-reflection diamond ATR accessory (DurasamplIR™, SensIR Technologies, incidence angle: 45°) was used to acquire spectra. Yeasts were cultivated in RPMI medium with (0.05 or 5 µg/mL) or without caspofungin for 16 h at 30°C with agitation. Cells were washed in PBS and one drop of yeast suspension in PBS was deposited on the ATR crystal. The PBS spectrum was used to remove the spectral background. The spectra of the fingerprint regions of interest were baseline-corrected at 1800 and 800 cm^-1^ and normalized to 1 in the region 1597 to 800 cm^-1^.

### Protein extraction and western blotting

Protein was extracted from 5 × 10^8^ cells cultivated in YPD in the absence or presence of caspofungin (5 μg/mL) for 3 h. After washing with sterile water, the pellet was resuspended in 250 μL of Laemmli buffer containing 0.5 μL of protease inhibitor (Cocktail Set III, Merck) and 50 μL of phosphatase inhibitor (Cocktail Set III, Merck). Cells were disrupted using the glass-bead method at 4°C, and the homogenate was heated to 100°C for 10 min. Crude cell extracts corresponding to 5 × 10^6^ cells per well were separated by SDS-PAGE (4-20%) for 45 min at 180 V. They were semi-dry transferred onto PVDF membranes (Bio-Rad). After incubation for 1 h in blocking solution (5% skimmed milk, PBS, 0.2% Tween 20), membranes were incubated for 12 h with the following primary antibodies: mouse anti-phospho-p38 MAP kinase (Thr180/182) 28B10 for the detection of Hog1p-P (Cell Signaling Technology) (dilution 1: 1000); mouse anti-phospho-p44/p42 MAPK (Thr202/Tyr204) for the detection of Cek1p-P and Mkc1p-P (Cell Signaling Technology) (dilution 1: 1500); and mouse anti-TAT1 monoclonal for the detection of tubulin as the loading control (ECACC #00020911) (dilution 1: 1000). After washing with 1 M NaCl and blocking solution, the membranes were incubated for 1 h at room temperature with the following secondary antibodies: ECL Plex anti-mouse Cy3 conjugate (GE Healthcare #PA43009V, 1: 5000) and ECL Plex anti-rabbit Cy5 conjugate (GE Healthcare #PA45011V, 1: 5000). After washing with blocking solution, the membranes were subjected to direct fluorescence detection using the ImageQuant™ LAS 4000 (GE Healthcare) according to the manufacturer’s recommendations.

### Macrophage cultivation and yeast infection

J774A.1 (ATCC TIB-67) murine macrophages were cultured in cRPMI medium (RPMI-1640 without phenol red, supplemented with 10% heat inactivated fetal bovine serum, 1 mM sodium pyruvate, and 2 g/L sodium bicarbonate), and infected as described previously (31). Briefly, J774 macrophages were adhered, overnight at 37°C in 5% CO2, in 96-well plates with clear well bottoms (Greiner Bio-one) at a concentration of 2 x 10^5^ macrophages per well in 200 µL of cRPMI. Three separate plates were set up to perform a time course analysis of the infection over 24 hours, at T30min, T5h and T24h. Yeast cells collected from overnight culture in YPD containing 5 μg/mL CFW, and supplemented or not with caspofungin at 0.05 or 5 μg/mL, and adjusted to a density of 1 × 10^6^/mL in cRPMI plus 5 5 μg/mL CFW, with or without caspofungin (0.05 or 5 μg/mL). J774 macrophages were infected with 200 µL of CFW-labeled yeasts treated or not with caspofungin at a MOI of 1 macrophage:1 yeast. As controls, PBS alone, macrophages stained with CFW, and yeasts stained with CFW were included in the plate. Caspofungin did not alter the growth or viability of the macrophages. For each *Candida* strain, macrophage infection was performed in triplicate and each condition was tested in quintuplet per experiment.

### Flow cytometry and fluorimetry

The macrophage mortality rate and the ratio of macrophages engaged in phagocytosis was measured using flow cytometry, as described previously (31). Briefly, macrophages were double-stained with anti-mouse CD16-APC (Beckman Coulter) and calcein-AM (Sigma-Aldrich) at 30 min, 5 h, and 24 h post-infection with CFW-labeled yeasts treated or not with caspofungin (0.05 or 5 μg/mL). Macrophage viability was calculated as the percentage of macrophages positive for both calcein-AM and anti-CD16-APC fluorescence in an infection assay compared to uninfected macrophages. Phagocytosing macrophages were quantified as the number of macrophages positive for calcein, anti-CD16, and CFW fluorescence. The fungal biomass ingested by the macrophages was evaluated by measuring the mean fluorescence intensity (MFI) of CFW-labeled yeasts per phagocytosing macrophage. The background of each fluorescent marker was removed using yeast cells alone labeled with anti-CD16-APC and calcein-AM, and macrophages alone labeled with CFW.

### Yeast survival rate

J774 cells were infected in triplicate with CFW-labeled yeasts treated or not with caspofungin. After co-incubation for 24 h, endocytosed yeast cells were released by lysing the J774 macrophages in 1 ml of 0.1% ice-cold Triton X-100 (Acros Organics). The yeast cells were counted using a Malassez hemocytometer and diluted to 1 x 10^3^ cells/ml in YPD. Then, 100 μl of yeast suspensions containing 100 cells were plated onto YPD in duplicates, incubated at 30°C for 24-48 h. CFU were counted, and the percentages of survival were determined.

## SUPPLEMENTAL MATERIAL

Supplemental data for this article may be found at:

## ACKNOWLEDGEMENTS

This work was supported by grants from the University of Bordeaux and the Centre National de la Recherche Scientifique (CNRS). The authors have no conflicts of interest to declare. We thank Merck Sharp & Dohme Corp. for providing caspofungin. We acknowledge the assistance of the Spectroscopy and Microscopy Service Facility of SMI LCPME (Université de Lorraine-CNRS; http://www.lcpme.ul.cnrs.fr/), and Nicolas Landrein for his help in designing figures.

